# Structures of TOG1 and TOG2 From the Human Microtubule Dynamics Regulator CLASP1

**DOI:** 10.1101/479766

**Authors:** Jonathan B. Leano, Kevin C. Slep

## Abstract

Tubulin-binding TOG domains are found arrayed in a number of proteins that regulate microtubule dynamics. While much is known about the structure and function of TOG domains in the XMAP215 microtubule polymerase family, less in known about the TOG domain array found in the CLASP family. The CLASP TOG array promotes microtubule pause, potentiates rescue, and limits catastrophe. How distinct the TOG domains of CLASP are from one another, from XMAP215 TOG domains, and whether they are positionally conserved across CLASP family members is poorly understood. We present the x-ray crystal structures of human CLASP1 TOG1 and TOG2. The structures of CLASP1 TOG1 and TOG2 are distinct from each other, from CLASP TOG3, and are positionally conserved across species. While studies have failed to detect CLASP TOG1 tubulin-binding activity, TOG1 is structurally similar to the free-tubulin binding TOG domains of XMAP215. In contrast, though CLASP TOG2 and TOG3 have tubulin binding activity, they are structurally distinct from the free-tubulin binding TOG domains of XMAP215. CLASP TOG2 has a convex architecture, predicted to engage a hyper-curved tubulin state. CLASP TOG3 has unique structural elements in the C-terminal half of its α-solenoid domain that modeling studies implicate in binding to laterally-associated tubulin subunits in the microtubule lattice in a mode similar to, yet distinct from XMAP215 TOG4. These findings highlight the structural diversity of TOG domains within the CLASP TOG array and provide a molecular foundation for understanding CLASP-dependent effects on microtubule dynamics.

## Introduction

Microtubules are highly dynamic, polarized eukaryotic cellular polymers [1–4]. Microtubules are composed of αβ-tubulin heterodimers that polymerize through lateral and longitudinal associations to form a cylindrical, polarized lattice with α-tubulin and β-tubulin exposed the microtubule minus and plus end respectively. Microtubule dynamics occur at both the plus and minus ends, but are primarily focused at the plus end. During phases of polymerization, tubulin heterodimers with GTP bound at the exchangeable site on β-tubulin incorporate into the lattice and define the “GTP cap”. Once incorporated into the lattice, the GTP in the exchangeable site is hydrolyzed to GDP. It is the structural transition of tubulin subunits in the microtubule lattice from a GTP-bound state to a GDP-bound state that underlies the polymer’s dynamic instability. Collectively, dynamic instability includes phases of polymerization, depolymerization, and pause, with the transition to depolymerization termed catastrophe, and the transition out of depolymerization termed rescue. While dynamic instability is inherent to microtubules it is highly regulated in space and time by a host of microtubule associated proteins (MAPs). A key subset of MAPs include microtubule plus end binding proteins that localize to polymerizing microtubule plus ends [5–8].

Many microtubule plus end binding proteins form a complex network of interactions both with each other and the microtubule polymer. A master plus end binding protein family is the end binding (EB) protein family (EB1, EB2, and EB3) that preferentially binds the post-hydrolysis GDP P_i_ microtubule state, best-mimicked by GTPγS-bound microtubules [9]. EB members use a dimerization domain to recruit SxIP or LxxPTPh motif-containing proteins to the microtubule plus end [7,10–12]. Two prime factors that bind tubulin and are recruited to microtubule plus ends by EB1 (either directly or indirectly) are Cytosolic Linker-Associated Protein (CLASP) and ch-TOG [13–17]. Both ch-TOG and CLASP are critical for proper interphase microtubule dynamics as well as mitotic spindle structure and dynamics [13,18–20]. While ch-TOG promotes microtubule polymerization, CLASP promotes microtubule pause and rescue and limits catastrophe [13,21–26]. Mutations in CLASP family members result in aberrant microtubule dynamics that manifest in phenotypes ranging from abnormal mitotic spindle structure to defects in axon guidance [20,27–29]. How CLASP and ch-TOG mechanistically regulate microtubule dynamics is poorly understood.

While ch-TOG and CLASP differentially affect microtubule dynamics, they both employ an array of tubulin-binding TOG domains to regulate the microtubule polymer [13,19,30,31]. TOG domain structures were first determined from ch-TOG family members, revealing a 220-250 residue α-solenoid comprising six HEAT repeats (HRs) (A through F) that form a paddle-like structure [30,32]. The intra-HEAT loops that line one face of the TOG domain are highly conserved and are used to engage the tubulin heterodimer [30,32]. Structural work involving TOG1 and TOG2 from the *Saccharomyces cerevisiae* ch-TOG family member Stu2 demonstrated that TOG domain HRs A-D and HRs E-F engage regions of β- and α-tubulin respectively that are exposed on the cytosolic surface of the microtubule [33,34]. Elucidating the structural determinants that underlie ch-TOG family TOG architecture led to the prediction and subsequent confirmation that CLASP also contains an array of cryptic TOG domains that underlies its regulatory action on microtubule dynamics [30,31]. While human ch-TOG contains an N-terminal pentameric TOG domain array, CLASP1 contains three TOG domains (TOG1-3) followed by a C-terminal CLIP-170 interaction domain (CLIP-ID) (Fig 1A) [13,19,35]. TOG structures determined to date from ch-TOG and CLASP family members show dramatically different curvatures along the TOG domain’s α-solenoid axis that predict distinct interactions with tubulin [22,23,30–39]. Of note, the structure of CLASP1 TOG2 revealed a unique bent architecture that would require a significant conformational change in either the TOG domain and/or tubulin to enable each component to fully engage [31]. While TOG domains have diverse architectures, structural data collected to date indicates that distinct architectures are conserved within an array and play position-specific roles in the regulation of microtubule dynamics [35]. This has led to the hypothesis that ch-TOG and CLASP families use a common TOG array-based paradigm to regulate microtubule dynamics, but employ distinct TOG architectures along their respective arrays to differentially regulate the polymer’s dynamics.

**Fig 1.**
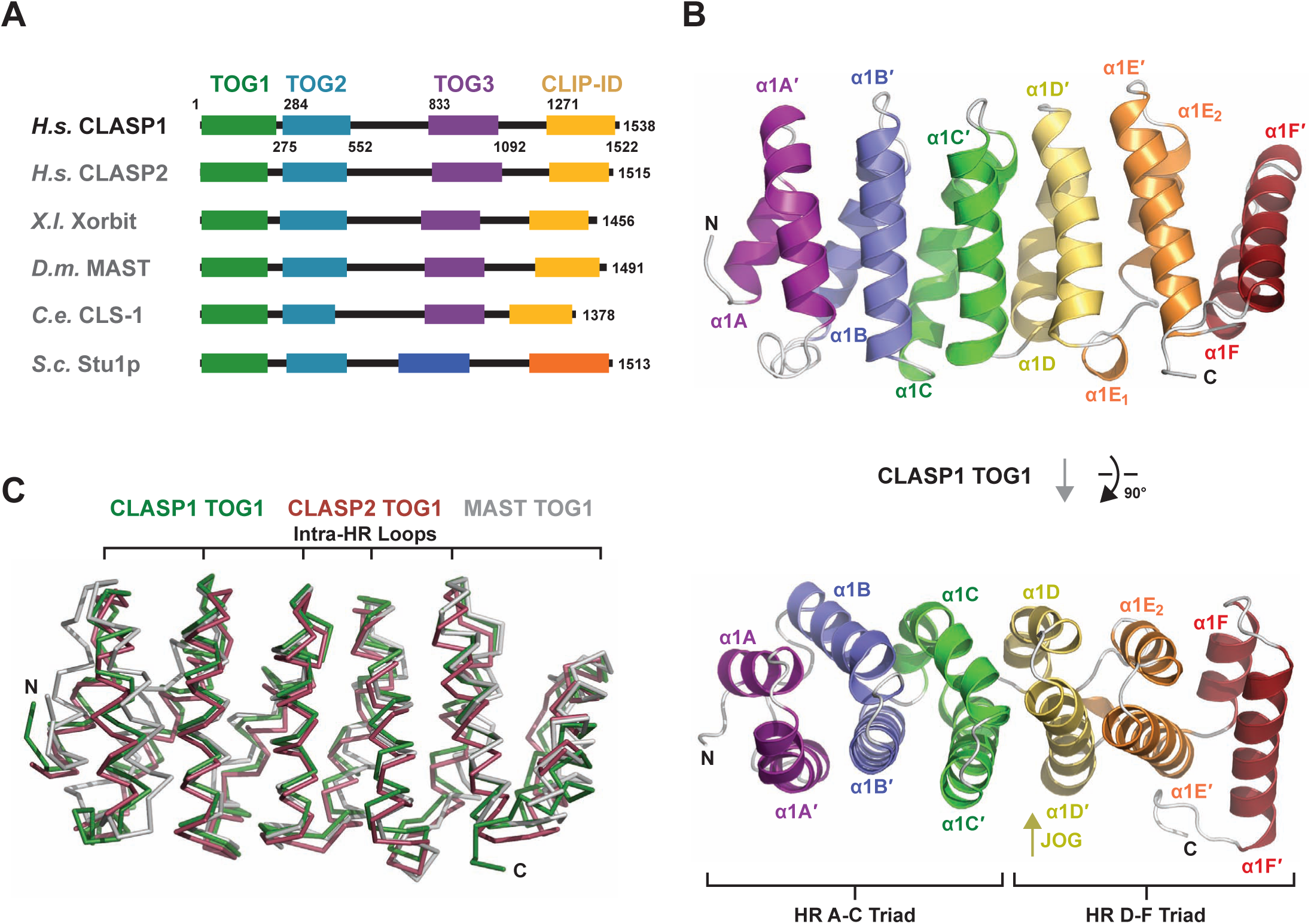
CLASP1 TOG1 forms a structurally conserved α-solenoid composed of six HRs. (A) Domain architecture of CLASP family members. *H. sapiens* (*H.s.*) CLASP1 and CLASP2, *X. laevis* (*X.l.*) Xorbit, *D. melanogaster* (*D.m.*) MAST, *C. elegans* (*C.e.*) CLS-1, and *S. cerevisiae* (*S.c.*) Stu1p. Stu1p’s two C-terminal domains are uniquely colored based on lack of homology to TOG3 and the CLIP-170 interaction domain (CLIP-ID) of other CLASP family members. (B) Architecture of *H. sapeins* CLASP1 TOG1 shown in cartoon format. The helices of each of the six HRs (A-F) are colored across the spectrum. The image at top is rotated 90° about the x-axis to present the view shown at bottom which focuses on the intra-HR loops. The HR α-solenoid is approximately 65 Å long. A translational shift between HR C and HR D prevents an overall supercoiling, common to α-solenoid domains, thereby facilitating an overall flat, paddle-like architecture for the TOG domain. (C) Structural alignment of CLASP1 TOG1 with *H. sapeins* CLASP2 TOG1 and *D. melanogaster* MAST TOG1 (PDB accession codes 5NR4 and 4G3A respectively, [22, 38]). Pairwise alignment using the Dali server [56] yields 1.2 Å rmsd across 220 Cα atoms for the CLASP1-CLASP2 TOG1 comparison, and 2.4 Å rmsd across 224 Cα atoms for the CLASP1-MAST TOG1 comparison.

Structural studies of CLASP family members to date have presented the structures of *Drosophila melanogaster* MAST TOG1, human CLASP2 TOG1, CLASP1 TOG2, CLASP2 TOG2, *S. cerevisiae* Stu1 TOG2, and mouse CLASP2 TOG3 [22,23,31,38,39]. *D. melanogaster* MAST TOG1 has a flat TOG architecture similar to Stu2 TOG1 [38]. Structures of human TOG2 from CLASP1 and CLASP2 are similar to one another and exhibit the bent architecture described above [31,39]. The structure of mouse CLASP2 TOG3 reveals a bent architecture, albeit not in the plane observed in CLASP TOG2, but perpendicular to this, such that the TOG3 HR D-F triad likely engages unique determinants on α-tubulin [39]. A similar, yet distinct, orthogonally bent architecture was observed in structures of TOG4 from ch-TOG family members [36]. These structural findings indicate that TOG domains in the CLASP family array each have distinct architectures and likely play distinct roles in tubulin-binding, microtubule affinity, and effects on microtubule dynamic instability. In support, recent studies of CLASP family members have implicated CLASP TOG2 as necessary and sufficient to limit microtubule catastrophe, and ascribes microtubule rescue activity to TOG3 [22,23]. While significant gains have been made in elucidating CLASP TOG structures, additional TOG structures from distinct family members are required to determine if these distinct TOG domain architectures are positionally conserved along the array.

Here we structurally characterize the first two TOG domains of human CLASP1. We present the X-ray crystal structure of CLASP1 TOG1 as well as a high-resolution structure of CLASP1 TOG2 (relative to our previously reported CLASP TOG2 structure [31]). These structures demonstrate that TOG architectures are positionally conserved across the CLASP family, but have non-equivalent architectures along the array. While tubulin-binding activity has not been detected for CLASP TOG1, CLASP1 TOG1 does conform to a tubulin-binding TOG architecture as observed in the structures of Stu2 TOG1 and TOG2 in complex with tubulin [33,34,40]. In contrast, CLASP1 TOG2, while containing Stu2 TOG-like tubulin binding determinants, adheres to a convex architecture across its tubulin-binding surface that predicts a unique tubulin-binding mode. The structures of CLASP1 TOG1 and CLASP1 TOG2 are architecturally distinct from one another as well as from the previously reported structure of CLASP2 TOG3. Modeling analyses suggest that TOG2 and TOG3 each engages tubulin in the microtubule lattice in a distinct fashion. This work highlights the emerging paradigm of a structurally diverse TOG domain array in which architecturally distinct domains each play unique roles in regulating microtubule dynamics.

## Materials and methods

### Protein expression and purification

Human CLASP1 TOG1 (residues 1-257) and TOG2 (residues 284-552) bacterial expression constructs were generated using the polymerase chain reaction method and individually sub-cloned into pET28 (Millipore Sigma, Burlington, MA). TOG1 and TOG2 construct expression and purification protocols were identical except as noted for the growth of TOG1 in minimal media containing selenomethionine, ion exchange chromatography, and final exchange buffer. Constructs were transfected into *Escherichia coli* (TOG1: B834 cells; TOG2: BL21 DE3 pLysS cells), grown to an optical density at 600 nm of 1.0 in media (TOG1: minimal media supplemented with seleno-L-methionine as described [41]; TOG2: Luria Broth) containing 50 µg/l kanamycin, the temperature lowered to 18° C, and protein expression induced with 100 µM Isopropyl β-D-1-thiogalactopyranoside for 16 hours. Cells were harvested by centrifugation, resuspended in buffer A (25 mM Tris pH 8.0, 200 mM NaCl, 10 mM imidazole, 0.1% β-ME) at 4° C, and lysed by sonication. Phenylmethylsulfonyl fluoride was added to 1 mM final concentration. Cells debris was pelleted by centrifugation at 23,000 x g for 45 minutes and the supernatant loaded onto a 5 ml Ni^2+^-NTA column (Qiagen, Hilden, Germany). The column was washed with 500 ml buffer A and protein eluted over a 250 ml linear gradient from 100% buffer A to 100% buffer B (buffer B = buffer A supplemented with 290 mM imidazole). Peak fractions were pooled, CaCl_2_ added to 1 mM final concentration, and 0.1 mg bovine α-thrombin added to proteolytically cleave off the N-terminal His_6_ tag on each construct. After a 24 hour incubation period at 4° C, protein was filtered over 0.5 ml of benzamadine sepharose (GE Healthcare Bio-Sciences, Pittsburgh, PA) and concentrated in a Millipore 10k MWCO centrifugal concentrator (Millipore Sigma, Burlington, MA). TOG1 was diluted into 100 ml buffer C (25 mM Tris pH 8.0, 0.1 % β-ME), and loaded onto a 10 ml Q-sepharose Fast Flow column (GE Healthcare Bio-Sciences, Pittsburgh, PA). Protein was washed with 200 ml buffer C and eluted using a 250 ml linear gradient between 100% buffer C and 100% buffer D (buffer D = buffer C + 1 M NaCl). Peak fractions were pooled and protein was concentrated and exchanged into storage buffer (10 mM Tris pH 8.0, 150 mM NaCl, 0.1% β-ME). TOG2 was diluted into 100 ml buffer E (25 mM Hepes pH 7.0, 0.1 % β-ME), and loaded onto a 10 ml SP-sepharose Fast Flow column (GE Healthcare Bio-Sciences, Pittsburgh, PA). Protein was washed with 200 ml buffer E and eluted using a 250 ml linear gradient between 100% buffer E and 100% buffer F (buffer F = buffer E + 1 M NaCl). Peak fractions were pooled and protein was concentrated and exchanged into storage buffer (25 mM Hepes pH 7.0, 200 mM NaCl, 0.1% β-ME).

### Crystallization, data collection, and structure determination

Selenomethionine-substituted TOG1 was crystallized via hanging drop: 2 µl of 10 mg/ml protein plus 2 µl of a 1 ml well solution containing 0.1 M 2-(*N*-morpholino)ethanesulfonic acid (MES) pH 6.5, 30% PEG 600, 10% glycerol, 18 °C. Crystals were grown from microseeds that originally crystallized in 1.5 M sodium malonate, pH 6.25, 18 °C. TOG2 was crystallized via hanging drop: 2 µl of 10 mg/ml protein plus 2 µl of a 1 ml well solution containing 22% PEG 3350 and 200 mM sodium citrate (pH 8.25), 18 °C. TOG1 and TOG2 crystals were frozen in paratone-N (Hampton Research, Aliso Viejo, CA) and diffraction data sets collected on single crystals at the Advanced Photon Source 22-ID beamline at 100 K. Data were processed using HKL2000 [42]. Attempts to determine phases for the TOG1 structure using single wavelength anomalous dispersion (SAD) phasing methods failed from crystals grown in 1.5 M sodium malonate, pH 6.25. Attempts to obtain phasing via molecular replacement (TOG1 search model: *Drosophila* MAST TOG1, PDB accession code 4G3A, chain A [38]) also failed. Thus, crystals grown in 0.1 M MES pH 6.5, 30% PEG 600, 10% glycerol, seeded from the original crystals grown in sodium malonate, were used for anomalous diffraction experiments and provided the SAD phasing for structure determination. The TOG2 structure was determined via molecular replacement using a TOG2 search model: human CLASP1 TOG2, PDB accession code 4K92, chain A [31].

Initial models were built using AutoBuild (PHENIX) followed by reiterative buildings in Coot [43] and subsequent refinement runs using phenix.refine (PHENIX) [44]. Refinement runs used real space, simulated annealing refinement protocols (temperatures: 5,000 K start, 300 K final, 50 steps), and individual B-factor refinement, using a maximum-likelihood target. Atomic displacement parameters were calculated for the TOG2 structure. The final refinement runs produced an R_free_ value of 24.4% for the TOG1 structure and an R_free_ value of 22.1% for the TOG2 structure. The final TOG1 model includes residues 1-234 for chain A, residues 1-236 for chain B, and 201 water molecules. The final TOG2 model includes residues 295-538 for chains A and B and 678 water molecules. Data collection and refinement statistics are summarized in Table 1. Structure images were generated using the PyMOL Molecular Graphics System, version 1.5.0.5 (Schrödinger, LLC, New York, NY). Electrostatic calculations used the PyMOL plugin APBS [45]. Pairwise structure comparisons and root mean square displacement (rmsd) values were calculated using the Dali server [46].

**Table 1.** Data processing and refinement statistics.

| **Crystal** | **CLASP1 TOG1** | **CLASP1 TOG2** |
| --- | --- | --- |
| **Data Collection** |  |  |
| Wavelength (Å) | 0.97964 | 1.0000 |
| Space group | P2<sub>1</sub> | P2<sub>1</sub> |
| Cell dimensions: a,b,c (Å) | 41.0, 114.0, 99.9 | 51.6, 67.4, 81.0 |
| Resolution (Å) | 50.0-2.15 (2.23-2.15) | 50.0-1.78 (1.84-1.78) |
| # Reflections: Measured / Unique | 148,462 (11,001) / 41,883 (3,667) | 319,655 (19,398) / 51,928 (4,492) |
| Completeness (%) | 92.3 (81.2) | 98.1 (85.4) |
| Mean redundancy | 3.5 (3.0) | 6.8 (4.3) |
| <I/σI> | 9.6 (1.96) | 15.1 (2.8) |
| R<sub>sym</sub> | 0.155 (0.646) | 0.095 (0.278) |
| **Refinement** |  |  |
| Resolution (Å) | 32.95-2.15 (2.20-2.15) | 38.1-1.78 (1.83-1.78) |
| R/ R<sub>free</sub> (%) | 19.0 (24.4) / 24.4 (33.3) | 17.7 (29.5) / 22.1 (38.0) |
| # Reflections, R/R<sub>free</sub> | 21,034 (1768) / 1930 (159) | 48317 (3683) / 1874 (153) |
| Total atoms: Protein / Water | 3,868 / 183 | 3,920 / 659 |
| Stereochemical ideality (rmsd): Bonds / Angles (Å/°) | 0.011 / 1.38 | 0.008 / 1.05 |
| Mean B-factors (Å<sup>2</sup>): Overall / Protein / Water | 28.8 / 28.6 / 31.3 | 29.0 / 27.1 / 39.9 |
| Ramachandran Analysis: Favored / Allowed (%) | 98.1 / 1.7 | 99.2 / 0.8 |
| **PDB accession code** | **6MQ5** | **6MQ7** |

Values in parentheses indicate statistics for the highest-resolution shell.

## Results and discussion

### The α-solenoid HEAT repeat structure of CLASP1 TOG1

To determine the structure of human CLASP1 TOG1 we crystallized a selenomethionine-substituted construct embodying residues 1 to 257 and collected a 2.15 Å resolution single wavelength anomalous dispersion dataset. Crystals belong to the space group P2_1_ and contain two protomers in the asymmetric unit. The structure was refined to R and R_free_ values of 19.0% and 24.4% respectively. Data collection and refinement statistics are presented in Table 1.

CLASP TOG1 is an α-solenoid structure, consisting of six HRs designated HR A through F (Fig 1B). We delineate the helices of each HR α and α’, followed by the number of the TOG domain in the array and the letter of the HR to which the helix belongs. The HRs conform to a general TOG-domain architecture. The first HR triad (HRs A-C) has a right-handed supercoil. The second HR triad (HRs D-F) is translated relative to the axis of the first HR triad, introducing a jog in the supercoil that gives the domain a flat, paddle-like architecture, rather than an elongated spiral common to other α-solenoid structures. HRs D and E are oriented with a right-handed twist relative to one another, while HRs E and F are oriented with a left-handed twist (Fig 1B, lower panel). The architecture of CLASP1 TOG1 is similar to that of *Homo sapiencs* CLASP2 TOG1 and *D. melanogaster* MAST TOG1 [22, 38], with overall pairwise Cα rmsd values of 1.2 and 2.4 Å for the CLASP1-CLASP2 and CLASP1-MAST comparisons respectively (Fig 1C, calculated using the Dali server [46]). CLASP1 TOG1 aligns best to MAST TOG1 (the more divergent comparison) across the TOG face composed of intra-HR loops.

### TOG1 is highly conserved across the surface defined by intra-HR loops

To examine TOG1 surface residue conservation, we generated a sequence alignment involving *H. sapiens* CLASP1, *Xenopus laevis* Xorbit, *D. melanogaster* MAST, and *Caenorhabditis elegans* CLS-1. We contoured conservation at 100% identity (dark green), 100% similarity (light green), and 75% similarity (yellow) and mapped this scheme on the *H. sapiens* CLASP1 TOG1 structure (Fig 2A,B). The domain face formed by intra-HR loops displayed the highest degree of conservation (Fig 2B, upper right), with additional conservation mapping to the face formed by the α’ helices of each HR (Fig 2B, upper left). TOG domains of the XMAP215 family are known to engage tubulin using the intra-HR loop surface. While CLASP1 TOG1 has a subset of intra-HR residues that are positionally similar to those found in XMAP215 TOG tubulin binding determinants, there are also significant differences. Specifically, XMAP215 family TOG domains primarily contain a tryptophan in the HR A loop. The homologous position in CLASP1 TOG1 is a valine (V17) that is not conserved (Fig 2C,D); *D. melanogaster* MAST TOG1 and *C. elegans* CLS-1 TOG1 have a methionine and proline in the equivalent position respectively (Fig 2A). Interestingly, while XMAP215 family TOG domains are primarily conserved across the HR A-C intra-HEAT loops, CLASP TOG1 conservation is primarily focused along the surface formed by the HR C-E intra-HR loops. Collectively, CLASP1 TOG1 has a canonical TOG domain architecture, but its unique surface residue conservation pattern suggests a binding function distinct from XMAP215 family TOG domains.

**Fig 2.**
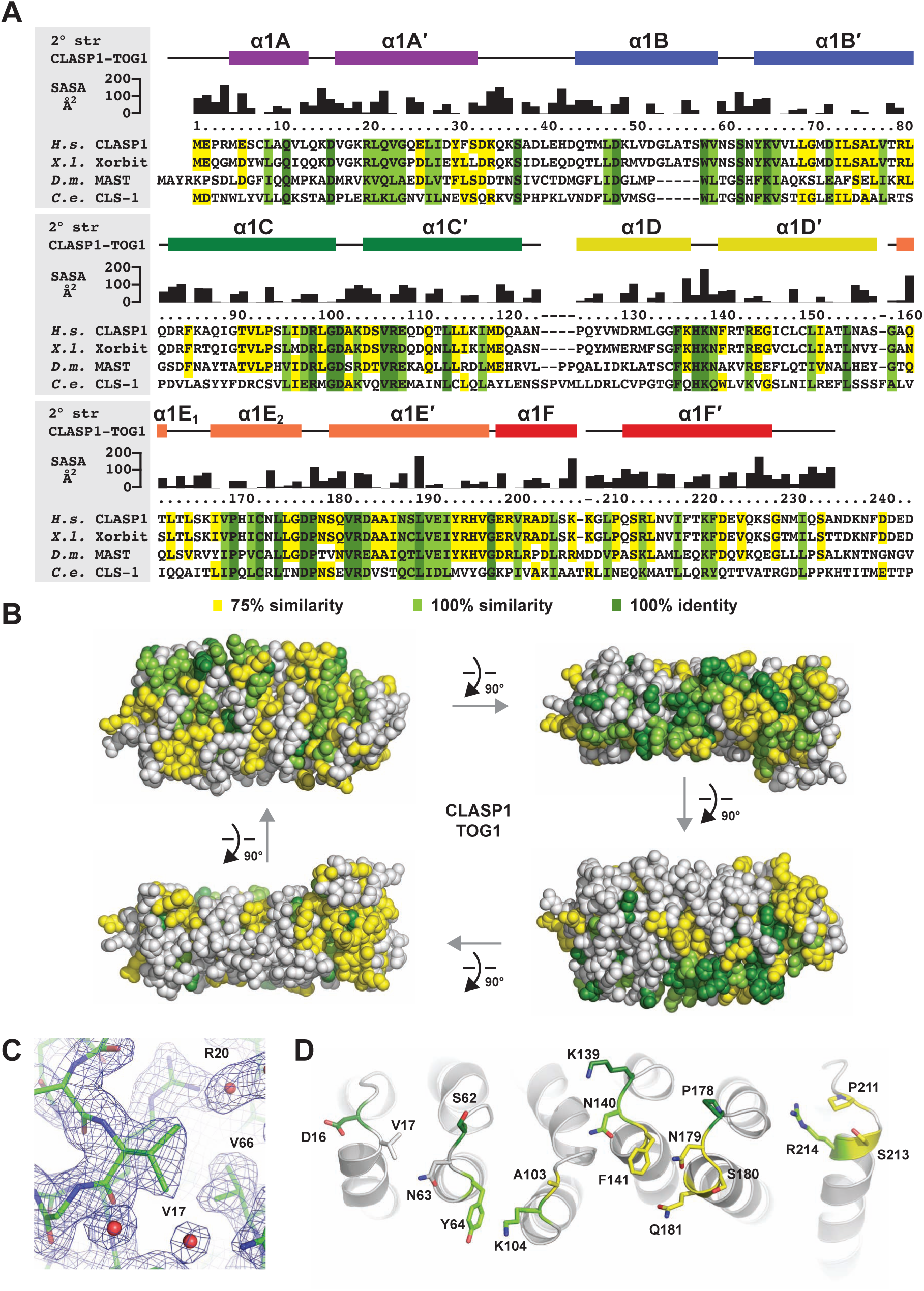
CLASP TOG1 has a conserved face delineated by intra-HR loops. (A) Sequence alignment of CLASP TOG1 from *H. sapiens (H.s.)* CLASP1, *X. laevis* (*X.l.)* Xorbit, *D. melanogaster (D.m.).* MAST, and *C. elegans (C.e.)* CLS-1. *H. sapiens* CLASP1 TOG1 2° structure and solvent accessible surface area (SASA) are shown above the alignment. Residues are highlighted based on 100% identity (dark green), 100 similarity (light green), and 75% similarity (yellow). (B) Cross-species conservation delineated in A, mapped on the CLASP1 TOG1 structure, and rotated in 90° steps about the x-axis. The orientation at upper left corresponds to the upper orientation in Figure 1B. Conservation primarily maps to the intra-HR loops that form the surface shown at upper right. Additional conservation maps to the face of the α-solenoid delineated by the α’ helix of each HR (face presented at upper left). (C) View of the HR A loop residue V17 shown in stick format with 2F_o_-F_c_ electron density shown in blue, contoured at 1.0 σ. (D) Intra-HR residues shown in stick format with conservation color-coded as in A and B.

### Mammalian CLASP TOG2 forms a conserved convex architecture

Previous structural work analyzing CLASP TOG2 revealed a highly bent, convex TOG architecture [31, 39]. To determine if this architecture was due to crystal packing, we determined the structure of CLASP1 TOG2 in a different space group, P2_1_, as compared to the initial structure that was determined in the space group P2_1_2_1_2_1_ [31]. Native diffraction data was collected to a resolution of 1.78 Å. The crystal contains two protomers in the asymmetric unit. The structure was solved by molecular replacement using the structure of *H. sapiens* CLASP1 TOG2 as a search model (PDB accession code 4K92 [31]). The structure was refined to R and R_free_ values of 17.7 % and 22.1% respectively. Data collection and refinement statistics are presented in Table 1.

The CLASP1 TOG2 structure determined in space group P2_1_ conforms to the bent TOG architecture observed in space group P2_1_2_1_2_1_ (Fig 3A). As previously observed, TOG2 has a bend between the HR A-C and HR D-F triads that orients the HR D-F intra-HR loop surface at ~30° relative to the plane established by the HR A-C intra-HR loops. This deviates significantly from the flat surface observed across XMAP215 family TOG intra-HR loops used to engage tubulin (discussed further below). In addition to its bent architecture, TOG2 has a conserved N-terminal helix, α2N, which is positioned alongside, and orthogonal to the α2B’ and α2C’ helices (Fig 3A). The two protomers in the P2_1_ asymmetric unit have an overall Cα rmsd of 1.4 Å. Comparing protomers determined in the P2_1_ space group to those determined in the P2_1_2_1_2_1_ space group showed little structure deviation with low overall Cα pairwise rmsd values that ranged from 0.3 to 1.3 Å. Comparison of the CLASP1 TOG2 protomers with the structure of CLASP2 TOG2, which has 81% sequence identity, yielded overall Cα rmsd values that ranged from 1.0-1.3 Å (Fig 3B). Collectively, the bent architecture of CLASP TOG2 domains is conserved and reflects a key structural state of the domain.

**Fig 3.**
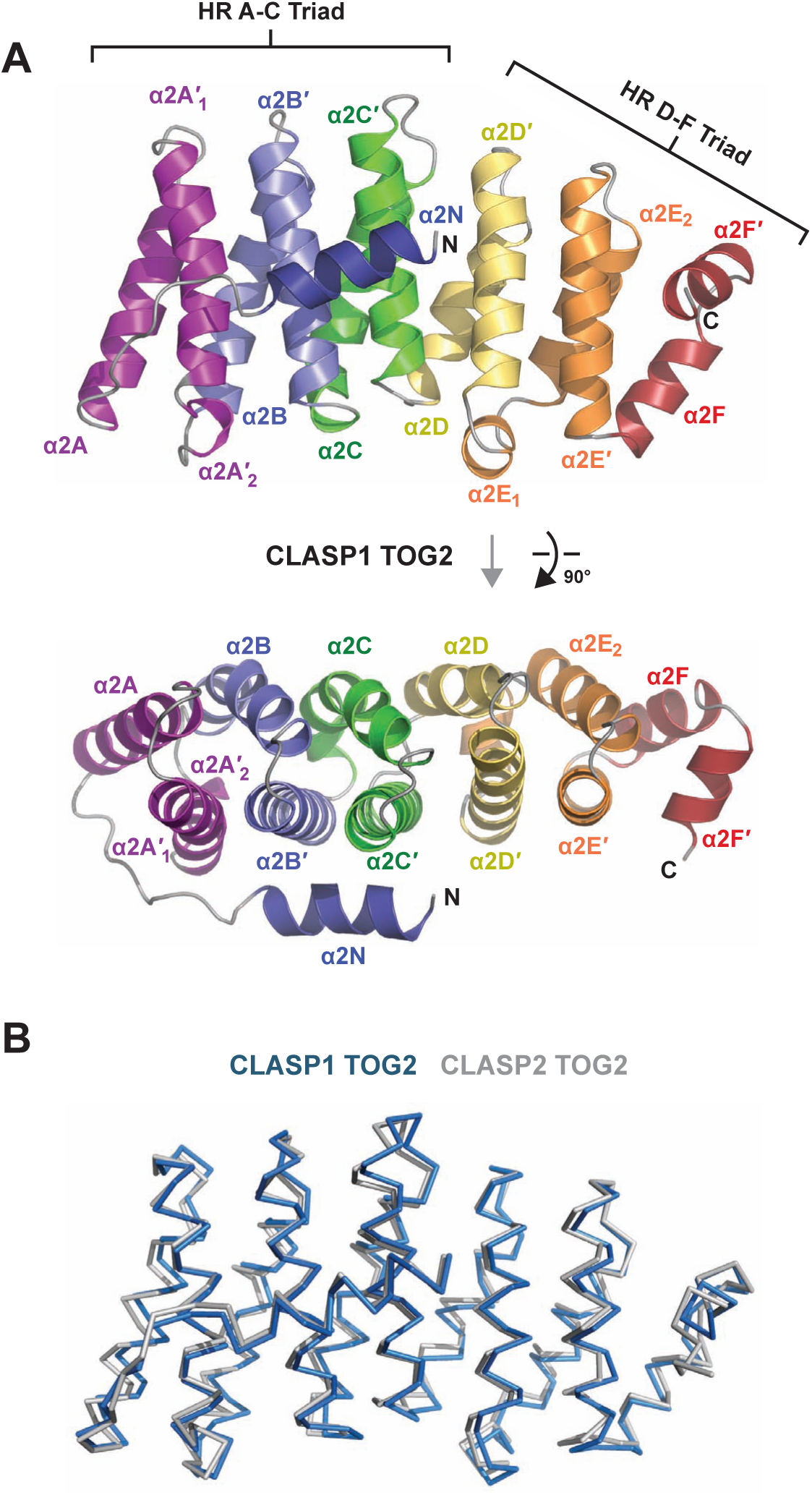
CLASP1 TOG2 forms a convex α-solenoid, structurally homologous to CLASP2 TOG2. (A) Architecture of *H. sapiens* CLASP1 TOG2 shown in cartoon format. The helices of each of the six HRs (A-F) are colored across the spectrum. CLASP1 TOG2 has a unique N-terminal helix, α2N, that runs orthogonal to and contacts the α’ helices of HRs B and C. The image at top is rotated 90° about the x-axis to present the view shown at bottom which focuses on the intra-HR loops. The HR α-solenoid is approximately 65 Å long. In contrast to canonical TOG structures determined to date, which form a relative straight surface across their intra-HR loops, CLASP TOG2 forms a bent, convex architecture across this surface (image at top). (C) Structural alignment of CLASP1 TOG1 and CLASP2 TOG2 (PDB accession code 3WOY, [39]). Pairwise alignment using the Dali server [56] yields 1.3 Å rmsd across 244 Cα atoms for the CLASP1-CLASP2 TOG2 comparison.

### TOG2 is highly conserved across the convex intra-HR loop surface

We next examined TOG2 surface residue conservation using the same species and conservation criteria laid forth in our TOG1 analysis (Fig 4A,B). As observed with TOG1, the domain face formed by intra-HEAT loops displayed the highest degree of conservation (Fig 4B, upper right), with additional conservation mapping to the orthogonally-positioned α2N helix (Fig 4B, upper left). A significant amount of surface residue conservation mapped to the remaining faces of the domain, pertaining primarily to the 75% similarity criteria. Of the highly conserved intra-HR loops, most of the conservation maps to the first triad, HR A-C. The HR A loop contains a conserved tryptophan, positioned equivalent to the conserved HR A loop tryptophan found in XMAP215 family TOG domains that engage β-tubulin (Fig 4C,D) [30,32–34]. A significant, yet lower degree of conservation maps to the second triad’s (HR D-F) intra-HR loops. Overall, TOG2 conservation implicates the intra-HR loops surface as a prime protein-protein interaction surface (implications for tubulin binding are discussed further below) while conservation across the remaining faces implicate these regions as likely protein-interaction surfaces as well. The degree of conservation across TOG2’s surfaces likely reflects the extent to which its cognate binding partners can co-evolve their interaction determinants. Specifically, the extent to which tubulin is conserved across species demands high cross-species conservation for the interaction determinants of its binding partners while additional factors that may bind other TOG2 surfaces (e.g. those involved in the auto-inhibition of TOG2 activity [22]) may be less evolutionarily constrained, permitting more variability and co-evolution of their respective binding interfaces.

**Fig 4.**
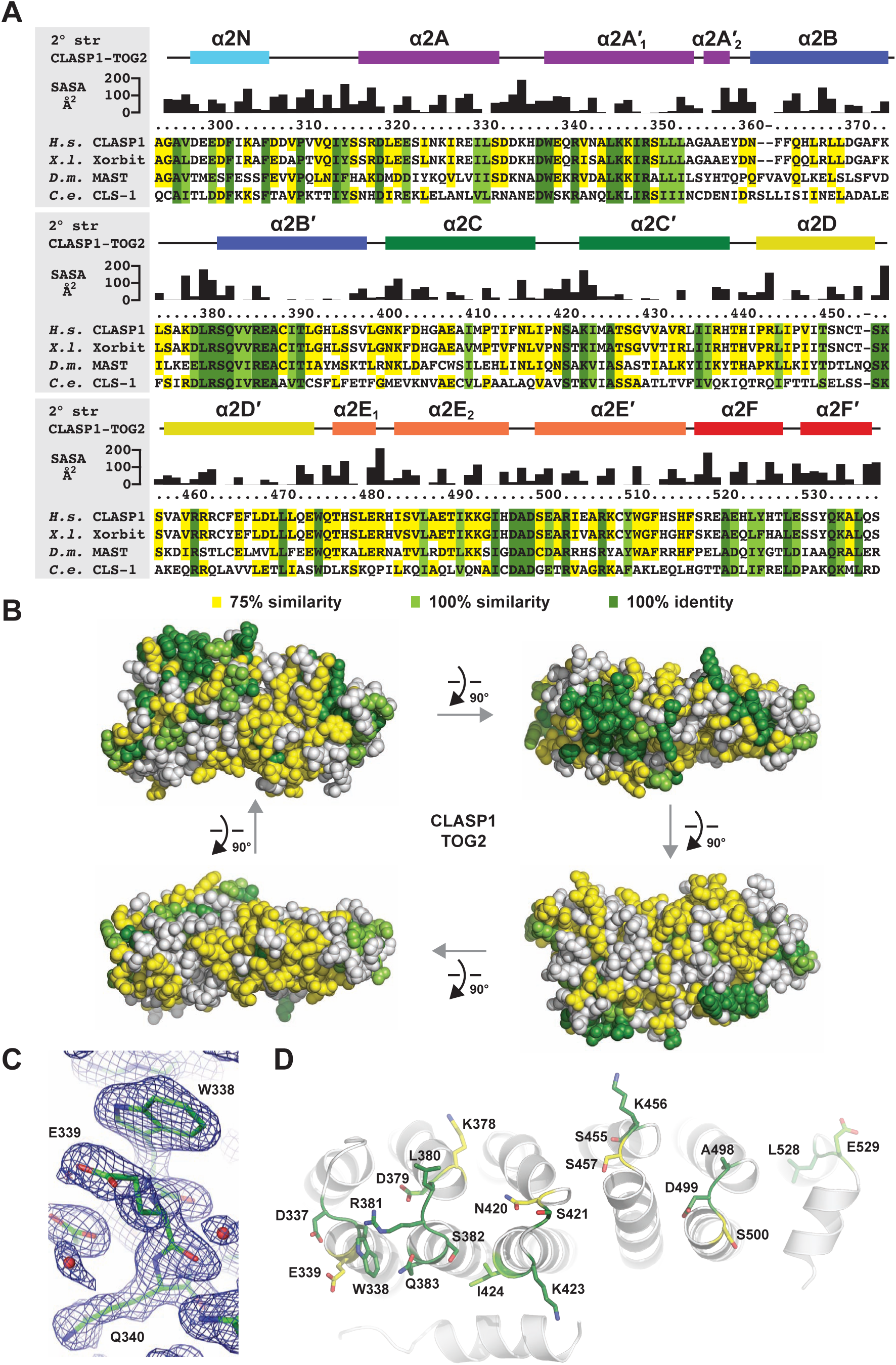
CLASP TOG2 has a conserved face delineated by intra-HR loops. (A) Sequence alignment of CLASP TOG2 from *H. sapiens (H.s.)* CLASP1, *X. laevis (X.l.)* Xorbit, *D. melanogaster (D.m.)* MAST, and *C. elegans (C.e.).* CLS-1. *H. sapiens* CLASP1 TOG2 2° structure and solvent accessible surface area (SASA) are shown above the alignment. Residues are highlighted based on 100% identity (dark green), 100% similarity (light green), and 75% similarity (yellow). (B) Cross-species conservation delineated in A, mapped on the CLASP1 TOG2 structure and rotated in 90° steps about the x-axis. The orientation at upper left corresponds to the upper orientation in Figure 3A. Conservation primarily maps to the intra-HR loops that form the surface shown at upper right. (C) View of the HR A loop residue W338 shown in stick format with 2F_o_-F_c_ electron density shown in blue, contoured at 1.0 σ. (D) Intra-HR residues shown in stick format with conservation color-coded as in A and B.

### The CLASP TOG array is structurally diverse

We next analyzed structural diversity across the CLASP TOG array, comparing our structures of CLASP1 TOG1, TOG2, as well as the previously reported structure of CLASP2 TOG3. We used the Dali server to structurally align the domains and calculate and overall rmsd value for corresponding Cα atoms [46]. Comparative analysis yielded the following high rmsd values: CLASP1 TOG1 versus CLASP1 TOG2: 3.3 Å rmsd; CLASP1 TOG1 versus CLASP2 TOG3: 2.9 Å rmsd; and CLASP1 TOG2 versus CLASP2 TOG3: 3.4 Å rmsd (Fig 5A). To determine whether specific subdomains contributed to this structural diversity, we again used the Dali server and analyzed the Cα rmsd across the TOG domains for each HR triad: HR A-C and HR D-F. The HR A-C triads aligned best, with rmsd values ranging from 1.9 Å (CLASP1 TOG1 versus CLASP2 TOG3) to 2.2 Å (CLASP1 TOG1 and CLASP2 TOG3 versus CLASP1 TOG2)(Fig 5A). In contrast, the HR D-F triads had a higher degree of structural variance, with rmsd values ranging from 2.3 Å (CLASP1 TOG1 versus CLASP1 TOG2) to 2.9 Å (CLASP1 TOG2 versus CLASP2 TOG3)(Fig 5A). Thus, while the first triad is structurally conserved across the CLASP TOG array, the second triad exhibits a higher degree of structural diversity. To determine if the relative positioning of the triads in each TOG domain also contributes to structural diversity across the CLASP TOG array, we structurally aligned the full TOG domains using the Cα coordinates from each domain’s respective HR A-C triad (Fig 5B). While TOG1 is relatively flat across the intra-HR loop surface, the alignment highlighted the relative bend between TOG2’s two triads that angles HR D-F downwards (Fig 5B, top panel). While TOG3 is flat across the HR A-E intra-HR loop surface, the HR F intra-HR loop region is positioned downward from the HR A-E intra-HR loop surface. When the intra-HR loop surfaces of TOG3 are viewed from above (Fig 5B, bottom panel), additional shifts (orthogonal to the relative bend observed in TOG2) are evident. While TOG1 bends to the side of the domain defined by the HR α’ helices, TOG2 is relatively straight. In contrast, TOG3 bends in the opposite direction, towards the side of the domain defined by the HR α helices. Collectively, the CLASP TOG array is structurally diverse, driven primarily by architectural diversity in the HR D-F triads as well as the relative positioning of the triads in the respective TOG domain. While each TOG domain in the array is architecturally distinct, the intra-HR loop surface of each domain is dominated by basic electrostatics and hydrophobic content (Fig 5C), suggesting that these conserved surfaces interact with the surface of a cognate partner that is negatively charged.

**Fig 5.**
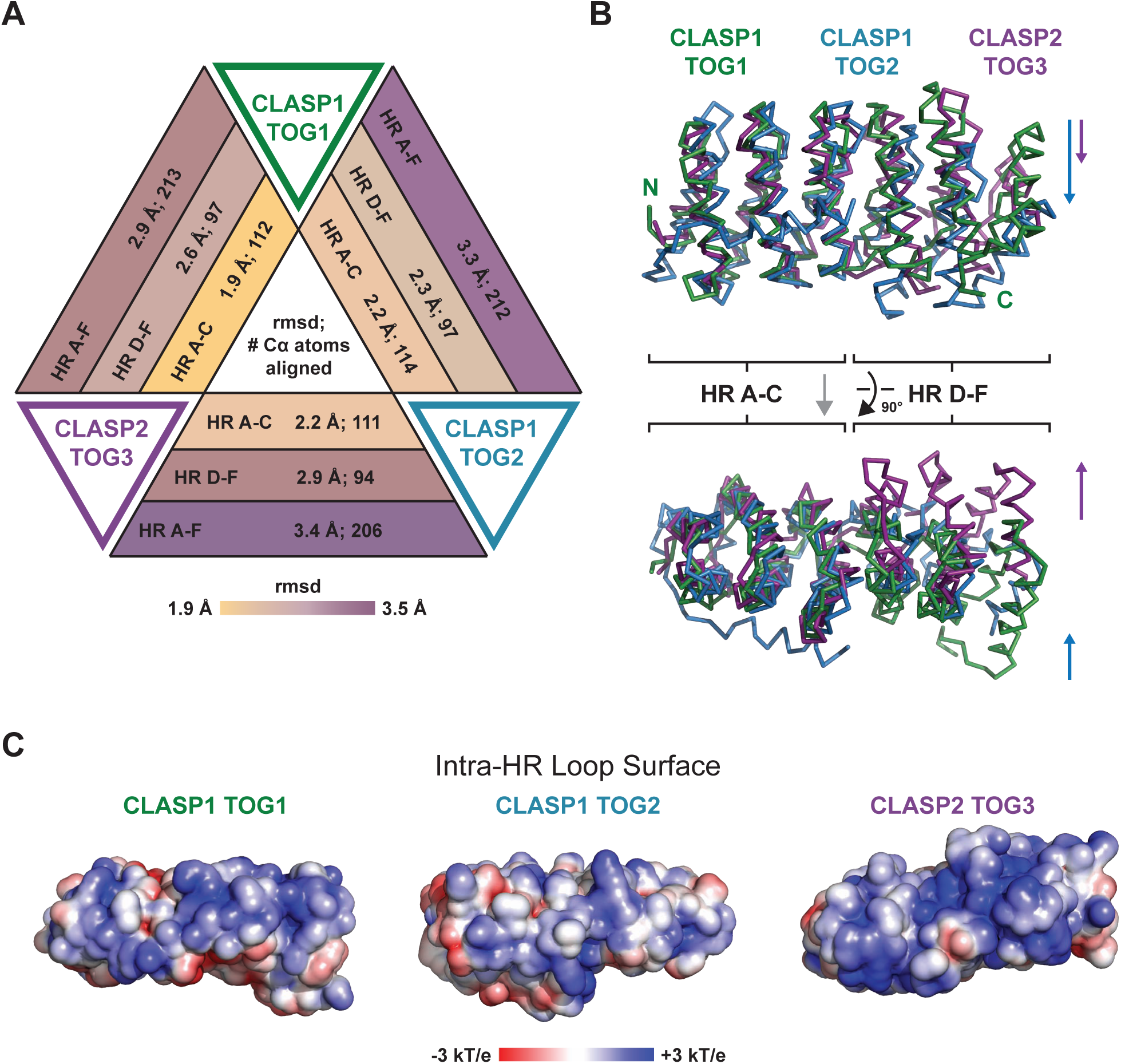
CLASP TOG1, TOG2, and TOG3 have unique domain architectures. (A) Comparison of CLASP1 TOG1, CLASP1 TOG2, and CLASP2 TOG3 (PDB accession code 3WOZ, [39]) structures using rmsd analysis of corresponding Cα atoms across the three domains (Dali server [56]). Pairwise analysis was performed, comparing HRs A-C, D-F, and all six HRs: A-F. The HR A-C triad exhibits a higher degree of pairwise structural homology with rmsd values ranging from 1.9-2.2 Å while the HR D-F triad has lower pairwise structural homology, with rmsd values ranging from 2.3-2.9 Å. (B) Structural comparison of CLASP TOGs 1-3. The TOG2 and TOG3 HR A-C triad was structurally aligned to the CLASP TOG1 HR A-C triad using the Dali server [56]. The alignment highlights the relative differential positioning of the HR D-F triads across the CLASP TOG array. Image at top is oriented with the intra-HR loops at the top of the domains. The image below was produced after a 90° rotation about the x-axis and focuses on the surface composed of intra-HR loops. (C) Electrostatic surface potential mapped on the structures of CLASP1 TOG1, TOG2, and CLASP2 TOG3. The surface of each domain presented is the surface composed of intra-HR loops, oriented as presented in the lower image of panel B.

### Varied CLASP TOG architectures suggest distinct tubulin binding modes

While structures of XMAP215 family TOG domains from the yeast member Stu2 bound to tubulin have informed how these TOG domains engage tubulin [33,34], how CLASP TOG domains bind tubulin remains unknown. Studies to date have implicated CLASP TOG2 and TOG3 in tubulin-binding, however, binding between TOG1 and tubulin (free or lattice bound) has not been detected [30,31,38–40]. To gain insight into CLASP TOG-tubulin binding modes and the potential basis for the lack of detectable TOG1-tubulin binding, we superimposed CLASP TOG domains on the Stu2 TOG2-tubulin structure and analyzed the modeled complexes. To generate these models, we aligned CLASP TOG domains to Stu2 TOG2 by superpositioning the first HR triads of each domain using the Dali server [46]. The basis for this was the structural conservation noted across the first HR A-C triad, which extended to Stu2 TOG2’s first triad. Stu2 TOG2 engages the curved state of the αβ-tubulin heterodimer, which reflects tubulin’s free state found in solution and at polymerizing and depolymerizing microtubule plus ends, as compared to the straight conformation found along the length of a microtubule [33,47,48]. Stu2 TOG2 HRs A-D engage β-tubulin while HRs E-F engage α-tubulin (Fig 6A) [33]. Key Stu2 TOG2-tubulin interaction determinants are a HR A loop tryptophan, an alanine and asparagine in the HR B loop, basic residues in the HR C, E, and F loops, and a threonine and proline residue in the HR D loop (Fig 6B). While there is no evidence that CLASP TOG1 interacts with tubulin, CLASP1 TOG1 superimposes surprisingly well on the Stu2 TOG2-αβ-tubulin structure, with all intra-HEAT loops assuming an interaction mode that complements the surface of the tubulin heterodimer with HR E intercalating the grove between α- and β-tubulin in a mode similar to Stu2 TOG2 HR E. Similarities between CLASP1 TOG1 and Stu2 TOG2 tubulin-binding determinants include the following: CLASP1 TOG1 has an asparagine in the HR B loop (N63), as well as basic residues in HR C, E, and F, that are positionally equivalent to Stu2 TOG2 K428, R519, and K549. While CLASP1 TOG1 does not have a basic residue in HR C equivalent to K427, it does have an arginine in HR D (R142), the guanidinium group of which is positioned to occupy the same relative space as the Stu2 TOG2 K427 side chain amine. CLASP1 TOG1 also has residues that are distinct from Stu2 TOG2 tubulin-binding determinants: as noted above, CLASP1 TOG1 lacks the conserved HR A loop tryptophan commonly found in XMAP215 TOG domains. Instead, CLASP1 TOG1 has a valine (V17) at the equivalent position. Additional intra-HR residues that are distinct to CLASP TOG1 are highlighted by conserved determinants in the HR D-F triad including HR D K139 and F141, HR E P178, and HR F S213 and R214 (Figs 2D and 6B). Overall, CLASP1 TOG1 has an architecture that aligns well with the tubulin-binding TOG domains of Stu2, some residues are similar in nature to XMAP215 family tubulin-binding determinants, but others are unique to the CLASP family. Why CLASP1 TOG1 does not bind tubulin, and what the identity of its binding partner is remains to be determined. Recent studies have implicated TOG1 in non-tubulin binding roles including kinetochore localization and regulating CLASP-dependent effects on microtubule dynamics by inhibiting the activity of TOG2 [22, 40].

**Fig 6.**
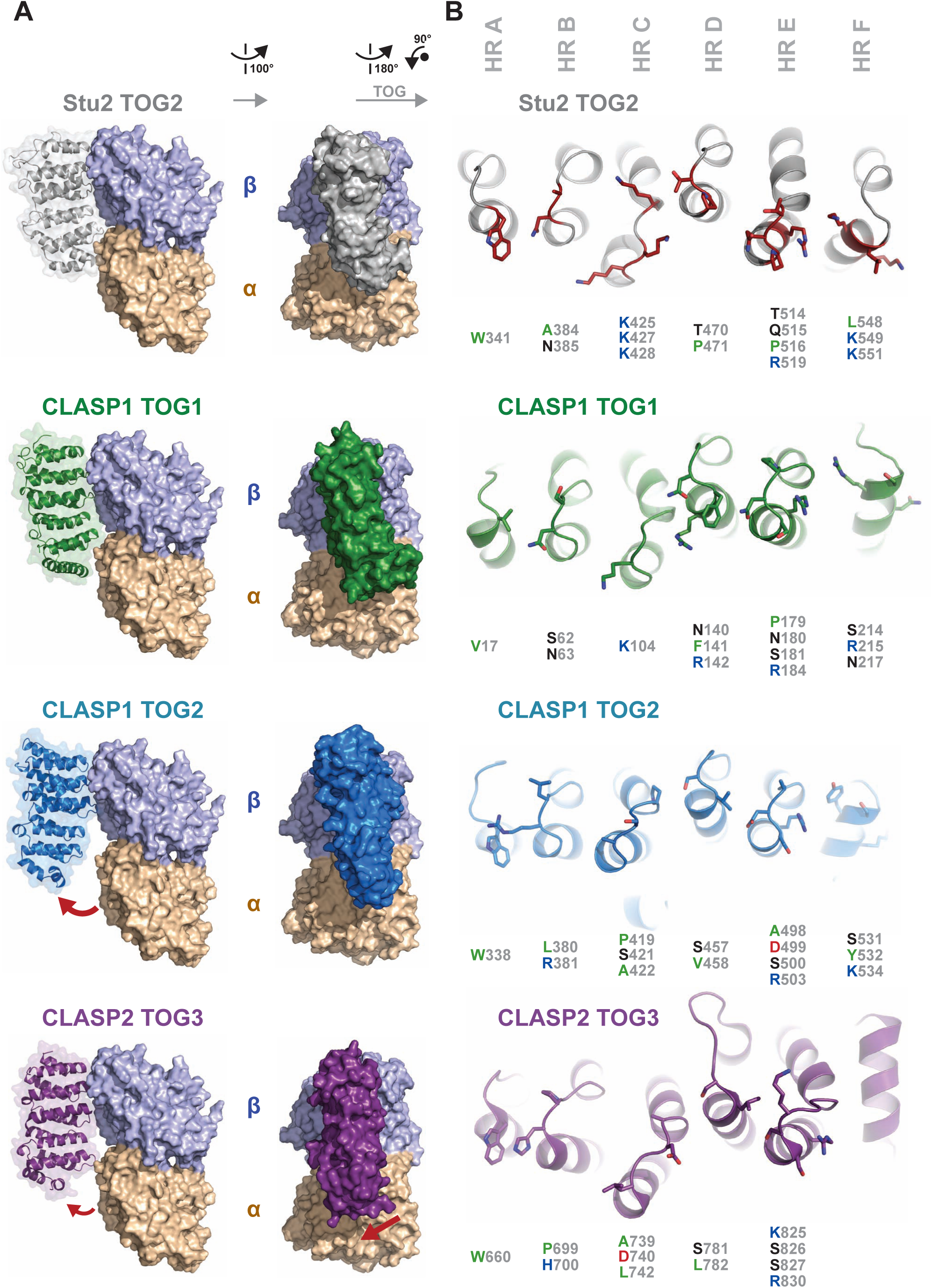
Distinct CLASP TOG architectures predict distinct tubulin binding properties. (A) CLASP TOG domains modeled on tubulin based on the structure of the XMAP215 microtubule polymerase family member Stu2 TOG2 in complex with αβ-tubulin (PDB accession code 4U3J [34], shown at top; Stu2 TOG2 in grey, α- and β-tubulin shown in sand and lavender respectively). Stu2 TOG2 intra-HR loop residues are involved in αβ-tubulin binding. HRs A-D engage β-tubulin while HRs E-F engage α-tubulin. To generate models of CLASP TOG domains bound to tubulin in a similar mode, the first HR triad (A-C) from each of CLASP’s TOG domains was structurally aligned to the Stu2 TOG2 HR A-C triad using the Dali server [56]. TOG domains at left are shown in cartoon format along with a transparent molecular envelope. The images at right were generated after a 90° rotation about the y-axis and depict each TOG domain in surface representation. The CLASP2 TOG3 structure is from PDB accession code 3WOZ [39]. (B) Comparative analysis of the intra-HR loop residues of each of CLASP’s TOG domains, which in Stu2 TOG2 are used to bind tubulin. The orientation of each domain, relative to the orientation shown in A (right panel) was generated after a 180° rotation about the y-axis, followed by a 90° counterclockwise rotation about the z-axis.

Unlike CLASP1 TOG1, the modeling of CLASP TOG2 and TOG3 onto tubulin using the Stu2 TOG2-tubulin complex as a guide yielded significant gaps between the second HR triad and αβ-tubulin (Fig 6A). For CLASP1 TOG2, the bent architecture of the domain angles HRs D-F away from αβ-tubulin. Interestingly, CLASP1 TOG2 has many Stu2 TOG2-like tubulin binding determinants across its intra-HR loops including HR A W338 (equivalent to Stu2 W341), HR D S457 and V458 (equivalent to Stu2 T470 and P471 respectively), HR E R503 (equivalent to Stu2 R519), and HR F K534 (equivalent to Stu2 R551)(Fig 6B). This suggests that CLASP TOG2 may engage tubulin across all of its intra-HR loops. If CLASP TOG2 adheres to a rigid conformation upon tubulin binding, this may in turn drive tubulin into a hyper-curved conformation that exceeds the curve observed in structures solved to date (e.g. see [33, 49]). Recent work has demonstrated that CLASP TOG2 is necessary and sufficient for CLASP-dependent microtubule anti-catastrophe activity [22, 23]. The unique bent architecture of CLASP TOG2 and its role limiting catastrophe may reflect the dynamic, curved protofilament architecture observed in both polymerizing and depolymerizing microtubules [48]. In contrast to the anti-catastrophe activity of CLASP TOG2, CLASP TOG3 has been found to promote microtubule rescue events [22]. Aligned with a distinct activity, our superpositioning model of TOG3 onto tubulin yields a distinct interaction mode (Fig 6A). CLASP TOG3 HR D is positioned away from β-tubulin and HR F is angled away from α-tubulin. The unique positioning of TOG3’s two HR triads relative to one another, and in comparison to the HR triads of Stu2 TOG2, leads to a unique positioning of the second HR triad on the surface of α-tubulin. While CLASP TOG3 has a unique TOG architecture, it retains a set of key intra-HR residues that are position equivalent to the tubulin-binding determinants of Stu2 TOG2 and are well oriented to engage αβ-tubulin. These include HR A loop W660 (equivalent to Stu2 W341), HR B P699 and H700 (equivalent to Stu2 A384 and N385 respectively), HR D S781 and L782 (equivalent to Stu2 T470 and P471 respectively), and HR E R830 (equivalent to Stu2 R519) (Fig 6B).

The distinct architecture of TOG3, modeled with a unique lateral shift on the tubulin heterodimer (Fig 6A) suggests that it may be involved in engaging a laterally associated tubulin subunit on the microtubule. To examine this, we superimposed the model generated of TOG3 bound to free tubulin (Fig 6A) onto the lattice coordinates of GMPCPP-bound tubulin (PDB accession code 3JAT [50]). As modeled, CLASP TOG3 makes contacts with the laterally-associated tubulin subunit on the adjacent protofilament. These contacts involve determinants in TOG3 HR B and the unique extended intra-HEAT loop of HR D (Fig 7A,B). The potential ability of CLASP TOG3 to bridge adjacent protofilaments may underlie its ability to promote rescue.

**Fig 7.**
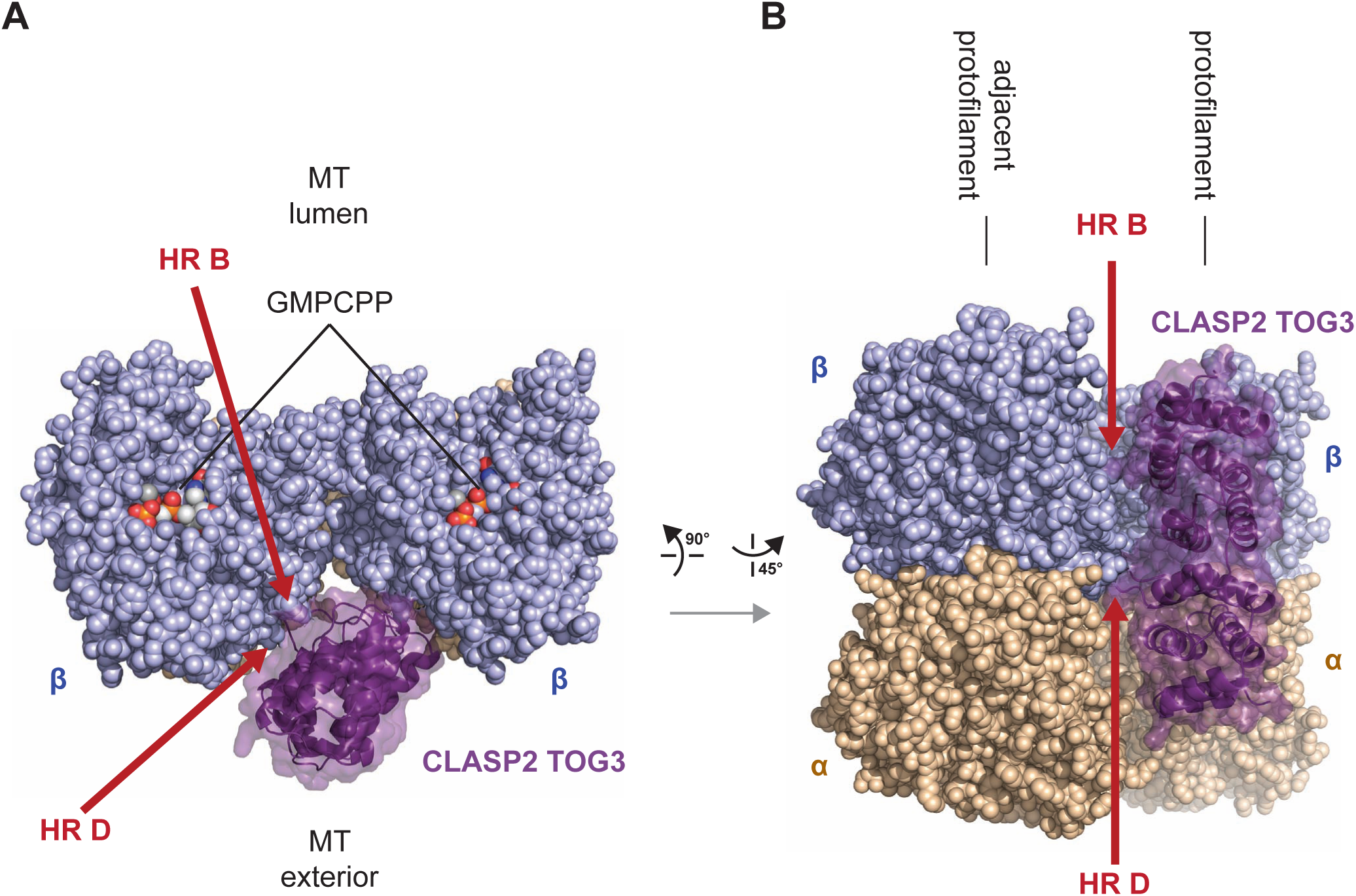
CLASP TOG3 is predicted to engage laterally associated tubulin on the microtubule lattice. (A) Model of CLASP2 TOG3 superpositioned on a microtubule. Shown are two laterally-associated tubulin heterodimers from neighboring protofilaments. TOG3 is shown in dark purple and bound to the tubulin heterodimer shown at right as in Fig 6A. The model generated of TOG3 bound to free tubulin (Fig 6) was superpositioned onto the lattice coordinates of GMPCPP-bound tubulin (PDB accession code 3JAT [50]). The tops of the β-tubulin subunits are shown (plus end oriented towards the viewer), looking into the bore of the microtubule with the luminal region oriented above and the microtubule exterior oriented below. CLASP TOG3 is shown bound to the tubulin subunit at right, and can engage the laterally associated tubulin subunit at left via intra-HR loop determinants in HR B and HR D. (B) Model as shown in (A), viewed from the microtubule exterior with the plus end oriented up. β-tubulin is shown in lavender, α-tubulin is shown in wheat. Potential TOG3 HR B and HR D contacts with the laterally-associated tubulin subunit are demarcated with red arrows.

## Conclusion

We have used crystallography to determine the structures of human CLASP1 TOG1 and TOG2. Our comparison of these structures to other CLASP TOG domain structures and to XMAP215 tubulin-binding TOG domains has highlighted a number of features of the CLASP TOG array: 1) each TOG domain along the CLASP TOG array has a unique architecture, 2) the structure of each specific TOG domain along the CLASP TOG array is well conserved across species, across paralogs, and across different crystal space groups, 3) the unique structures of these TOG domains and their respective conserved determinants correlates with each TOG domain having unique activities: TOG1 plays a non-tubulin binding role in kinetochore localization and relieving the auto-inhibition of TOG2’s anti-catastrophe activity, TOG2 plays a role in limiting catastrophe, and TOG3 plays a role in promoting rescue, 4) the unique structures of TOG2 and TOG3 predict distinct modes of tubulin binding on the microtubule lattice: TOG2 may preferentially engage a hyper-curved tubulin state and TOG3 may bridge adjacent tubulin subunits on neighboring protofilaments.

While our work highlights distinct structural features of the TOG array, a number of key questions remain outstanding. 1) What factor does TOG1 bind? While CLASP TOG1 has an architecture that is similar to the tubulin-binding architecture of Stu2 TOG2 bound to tubulin and has intra-HR determinants that are similar to those Stu2 TOG2 uses to bind tubulin, no tubulin-binding activity has been ascribed to CLASP TOG1. Instead of binding tubulin, CLASP TOG1’s conserved intra-HR loops may be involved in binding a kinetochore factor [40, 51], relieving the auto-inhibition of CLASP TOG2 activity [22], binding actin [52], or a yet to be determined factor. 2) What is the structural conformation of a TOG2- and a TOG3-tubluin complex as found on a microtubule? Will these structures yield insight into novel structural states of the tubulin heterodimer and will these states inform the mechanisms by which TOG2 and TOG2 affect microtubule dynamics? 3) While the CLASP TOG array is composed of structurally distinct TOG structures arranged in a specific order, can the array function properly if shuffled or do the unique architectures play synergistic position- and spatial-specific roles along the polarized microtubule lattice? 4) While the structural nature of CLASP’s three TOG domains (TOG1-3) has been elucidated, little is known about the C-terminal CLIP-ID [13]. The CLIP-ID is predicted to be composed of HRs and analysis suggests that it may conform to a TOG architecture as found in XMAP215 TOG5 [35]. Whether this is the case, whether the CLIP-ID has tubulin- or microtubule-binding activity, and how CLIP-170 binds the CLIP-ID remains to be determined.

Our work highlights the structural diversity of the known TOG domains that comprise the CLASP TOG array. This adds to our growing understanding of TOG domain structural diversity and the specific role these structures play when arrayed in different regulators of microtubule dynamics, including the CLASP family, the XMAP215 family of microtubule polymerases, and the Crescerin family that regulates microtubule dynamics in cilia [53–55].

## Accession numbers

**RSCB Protein Data Bank.** Coordinate and structure factors for the CLASP1 TOG1 and have been deposited with the accession code 6MQ5. Coordinate and structure factors for the CLASP1 TOG2 structure and have been deposited with the accession code 6MQ7.

## Acknowledgements

We thank the staff at the Argonne National Laboratory Advanced Photon Source SER-CAT beamlines 22-ID and 22-BM for support. Use of the Advanced Photon Source was supported by the U. S. Department of Energy, Office of Science, Office of Basic Energy Sciences, under Contract No. DE-AC02-06CH11357.

## Author Contributions

**Conceptualization:** Jonathan B. Leano, Kevin C. Slep

**Data Curation:** Jonathan B. Leano, Kevin C. Slep

**Formal Analysis:** Jonathan B. Leano, Kevin C. Slep

**Funding Acquisition:** Jonathan B. Leano, Kevin C. Slep

**Investigation:** Jonathan B. Leano, Kevin C. Slep

**Methodology:** Jonathan B. Leano, Kevin C. Slep

**Project Administration**: Kevin C. Slep

**Resources:** Kevin C. Slep

**Supervision:** Kevin C. Slep

**Validation:** Jonathan B. Leano, Kevin C. Slep

**Visualization:** Kevin C. Slep

**Writing – Original Draft Preparation:** Kevin C. Slep

**Writing – Review & Editing:** Jonathan B. Leano, Kevin C. Slep

